# Effect of Jasmonic Acid on Chlorophyll Content in Wheat (*Triticum aestivum* L.) Plants Infested with Russian Wheat Aphid (*Diuraphis noxia*)

**DOI:** 10.64898/2026.04.16.718935

**Authors:** Shanza, Mateen Ur Rehman, Farrah Farooq

## Abstract

Russian wheat aphid (*Diuraphis noxia* Homoptera; Aphididae) is a major pest that significantly reduces chlorophyll content and photosynthetic capacity in wheat (*Triticum aestivum* L.), leading to substantial crop yield losses. Jasmonic acid (JA) is a plant signaling molecule known to activate defense mechanisms against herbivorous insects. This study examined the effectiveness of exogenous jasmonic acid application in maintaining chlorophyll content during Russian wheat aphid infestation. A pot experiment was conducted with four treatments: control (no treatment), aphid infestation only, jasmonic acid application only, and jasmonic acid with aphid infestation. Results demonstrated that aphid infestation significantly reduced chlorophyll a (F = 42.565, P = 0.0001), chlorophyll b (F = 52.565, P = 0.0001), and total chlorophyll (F = 32.565, P = 0.0002) contents compared to healthy plants. Jasmonic acid treatment at 2 mM concentration effectively preserved all forms of chlorophyll, significantly counteracting aphid-induced chlorophyll depletion (P < 0.01). The protective effect of jasmonic acid was evident through the statistically significant interaction between aphid stress and JA application for all chlorophyll parameters. These findings suggest that foliar application of jasmonic acid can serve as an effective strategy to maintain photosynthetic capacity and plant vigor under Russian wheat aphid attack, thereby contributing to sustainable crop management and improved wheat production.

## Introduction

Wheat (*Triticum aestivum* L.), belonging to the family Poaceae, is one of the most important staple crops worldwide, providing approximately two-thirds of the global population with essential protein and calories [1]. The crop is grown in diverse geographical regions spanning from 45° south to 67° north latitude and has been cultivated for over 8,000 years in North Africa, Europe, and West Asia [2]. Despite its agricultural importance, wheat production faces considerable threats from biotic stresses, particularly insect pests [3].

Among these pests, the Russian wheat aphid (*Diuraphis noxia* Homoptera; Aphididae) represents one of the most destructive threats to cereal crop production globally [4]. This pest was first identified in 1901 and has since become a major problem in wheat-cultivating areas worldwide, including Pakistan, the United States, South Africa, and Mexico [5]. The aphid causes extensive damage through multiple mechanisms: direct feeding on phloem sap, injection of toxic substances through saliva that disrupts plant physiology, and transmission of viral pathogens [6]. These feeding mechanisms result in characteristic symptoms including chlorosis, leaf curling, stunted growth, necrosis, and ultimately, plant death [7].

The primary mechanism of Russian wheat aphid damage is the reduction of chlorophyll content in affected leaves [8]. By draining nutrients and water from the phloem and xylem tissues, the aphid disrupts photosynthetic processes, reducing the plant’s ability to produce energy necessary for growth and survival [9]. Poor chlorophyll concentration directly decreases photosynthesis and reduces the photochemical activity of the plant, potentially leading to complete cessation of plant growth [10]. This chlorophyll depletion represents a critical target for plant protection strategies, as maintaining photosynthetic capacity is essential for sustaining yield under pest pressure [11].

Traditional control methods for Russian wheat aphid, including chemical insecticides and pesticides, have been effective in reducing pest populations but are associated with significant environmental concerns, development of pesticide resistance in secondary pests, and toxicity to natural enemies [12]. These limitations have driven the search for sustainable and environmentally friendly pest management strategies [13]. Plant-based defense mechanisms, particularly those regulated by endogenous plant hormones, offer promising alternatives [14].

Jasmonic acid (JA), a naturally occurring plant hormone derived from linolenic acid, plays a central role in regulating plant defense responses to herbivory and pathogenic attack [15]. Unlike mechanical damage alone, herbivorous insect attack induces stronger physiological responses in plants through JA-mediated signaling pathways [16]. Jasmonic acid functions as a key signaling molecule that activates resistance genes, induces the synthesis of defensive compounds, attracts natural enemies of pests, and enhances antioxidant enzyme production [17]. Research has demonstrated that JA-induced resistance in wheat can reduce cereal aphid populations by up to 78% through natural parasitoids and predators, while also directly improving plant vigor and yield [18].

The exogenous application of jasmonic acid has been shown to enhance plant defense mechanisms at various developmental stages and under different stress conditions [19]. In wheat and tomato crops, JA application has been reported to induce significant resistance against aphid attack by catalyzing inhibitory enzymes such as proteases [20]. JA-treated plants produce volatile signaling compounds that attract natural predators, enhancing biological control. This multifaceted action makes jasmonic acid a promising candidate for integrated pest management [21].

Despite these promising reports, detailed quantitative studies on the direct effect of jasmonic acid application on chlorophyll preservation during Russian wheat aphid infestation in wheat remain limited [22]. Understanding whether and to what extent exogenous JA can maintain chlorophyll content under aphid attack is critical for developing practical management strategies [23]. This study was therefore designed to investigate the protective effect of jasmonic acid on chlorophyll content in wheat plants subjected to Russian wheat aphid infestation, with the hypothesis that JA application would preserve chlorophyll a, chlorophyll b, and total chlorophyll contents, thereby maintaining photosynthetic capacity under biotic stress.

## 2. Materials and Methods

### 2.1 Experimental Setup and Design

The experiment was conducted at the Integrated Genomics, Cellular, Developmental and Biotechnology Laboratory (IGCDBL) of UAF PARS campus. Eighteen plastic pots, each containing 10 kg of soil, were prepared and arranged in a randomized complete design (RCD) with three replicates per treatment. Wheat seeds (*Triticum aestivum* L.) were sown in each pot and allowed to germinate under controlled conditions with adequate moisture.

### 2.2 Treatment Application

Following an 8-week germination and growth period, plants were subjected to the following treatments:

1. Control: No treatment (healthy plants)
2. Aphid-infested control: Aphid infestation without jasmonic acid treatment
3. Jasmonic acid only: JA treatment without aphid infestation
4. Jasmonic acid + aphid: Combined JA treatment and aphid infestation

Jasmonic acid was prepared at a concentration of 2 mM by dissolving JA in distilled water. Plants designated for JA treatment received foliar sprays at 3-day intervals throughout the experimental period. Approximately 24 hours after the final JA application, plants were artificially infested with Russian wheat aphid (*Diuraphis noxia*) at standardized population densities. Untreated control plants received no spray application and were either left uninfested or infested with aphids.

### 2.3 Chlorophyll Extraction and Quantification

Fresh leaf tissue (0.5 g) was harvested from the upper portion of each plant and immediately ground using a mortar and pestle in 80% acetone until complete homogenization was achieved. The volume was adjusted to 5 mL with additional acetone, and the homogenate was centrifuged at 12,000 rpm for 15 minutes. The resulting filtrate was maintained at freezing temperature overnight to ensure complete precipitation of any residual cellular material.

Light absorbance of the clarified extract was measured using a Hitachi U-2001 spectrophotometer (Japan) at two wavelengths:

- 645 nm for chlorophyll b absorption
- 663 nm for chlorophyll a absorption

Chlorophyll concentrations were calculated using the extinction coefficients and formulas provided by Yoshida et al. (1976):

Chlorophyll b (mg/g): [22.9 × OD_{645} - 4.68 × OD_{663}] × V / (1000 × W)

Chlorophyll a (mg/g): [12.7 × OD_{663} - 2.69 × OD_{645}] × V / (1000 × W)

Total Chlorophyll (mg/g): [20.2 × OD_{645} + 8.02 × OD_{663}] × V / (1000 × W)

Where:

- V = total volume of acetone used in the extract (mL)
- W = fresh weight of leaf tissue (g)

### 2.4 Statistical Analysis

Data were analyzed by one-way analysis of variance (ANOVA) using COSTAT statistical software. Treatment means were compared, and P-values were determined at significance levels of P < 0.05 (*), P < 0*.*01 (), and P < 0*.*001 (*). The interaction between aphid stress and jasmonic acid treatment was also assessed through factorial ANOVA.

## 3. Results

### 3.1 Chlorophyll a Content

Analysis of variance revealed that chlorophyll a content differed significantly between healthy and aphid-infested wheat plants (F = 42.565, P = 0.0001). Healthy control plants demonstrated substantially higher chlorophyll a content compared to plants infested with Russian wheat aphids alone.

Jasmonic acid application had a significant positive effect on chlorophyll a maintenance (F = 2.396, P = 0.0062). Plants treated with 2 mM jasmonic acid maintained significantly higher chlorophyll a content compared to untreated controls, regardless of aphid infestation status. The most pronounced effect was observed in plants that received JA treatment prior to aphid infestation, demonstrating the protective capability of the hormone against chlorophyll degradation.

The interaction between aphid stress and jasmonic acid treatment was highly significant (F = 60.340, P = 0.0001), indicating that the protective effect of JA was specifically enhanced in the presence of aphid stress, suggesting an active defense mechanism activation.

**Table 1.**
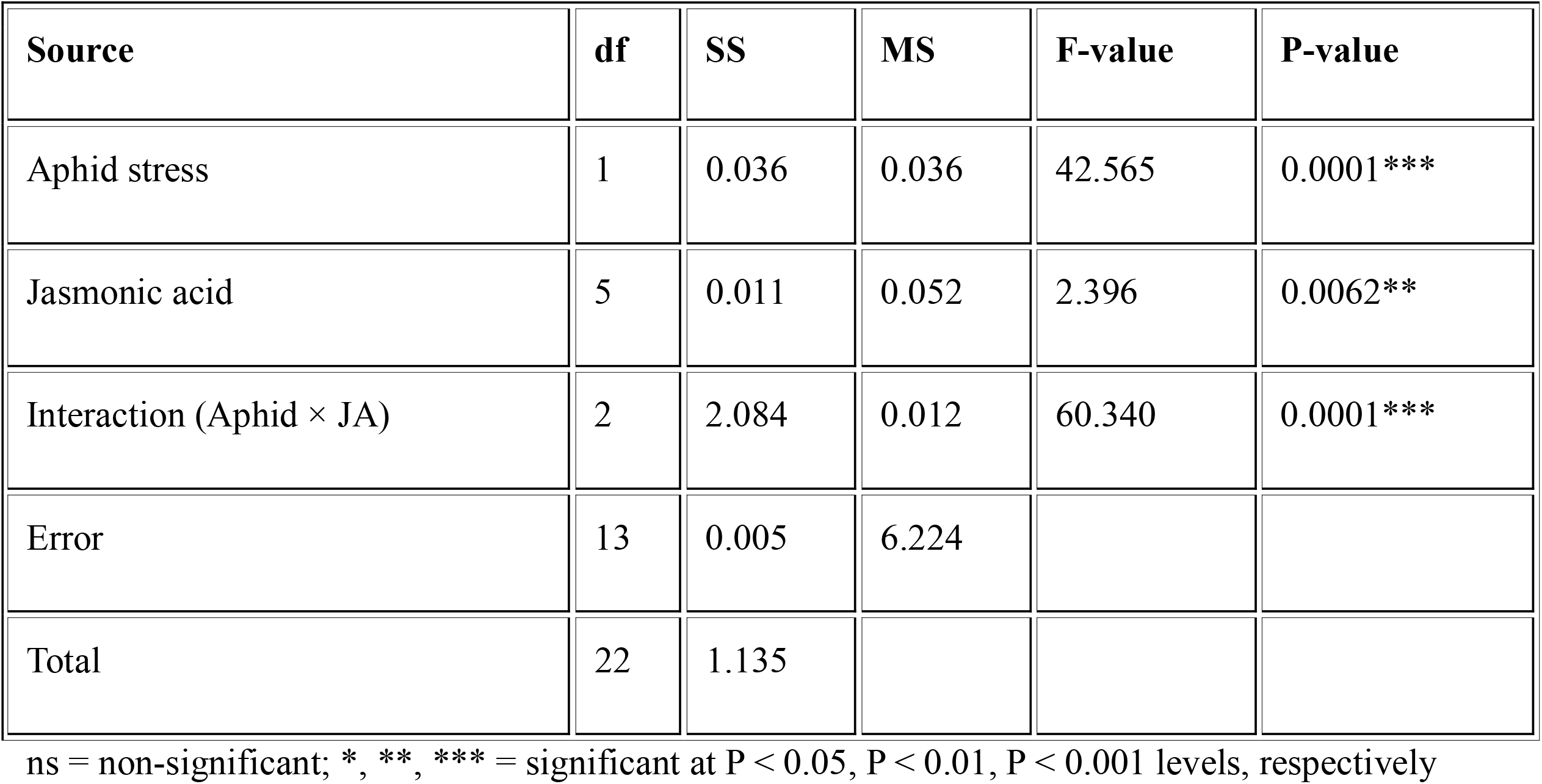
Analysis of Variance for Chlorophyll a Content.

### 3.2 Chlorophyll b Content

Chlorophyll b content also differed significantly between healthy and infested plants (F = 52.565, P = 0.0001), with aphid-infested plants showing marked reduction in chlorophyll b compared to healthy controls. This suggests that aphid infestation specifically targets photosynthetic pigment synthesis or accelerates pigment degradation.

Jasmonic acid treatment significantly increased chlorophyll b content (F = 8.396, P = 0.0063). The protective effect of 2 mM JA was evident in both aphid-free and aphid-infested plants, with particularly pronounced preservation of chlorophyll b in JA-treated plants subjected to aphid stress. This indicates that JA-mediated defense mechanisms are effective at preserving the accessory photosynthetic pigments necessary for light harvesting and energy transfer.

The interaction effect was highly significant (F = 60.740, P = 0.0002), demonstrating that jasmonic acid’s protective effect is amplified under aphid-induced stress conditions, suggesting coordinated activation of defense and photosynthetic maintenance pathways.

**Table 2.** Analysis of Variance for Chlorophyll b Content.

| Source | df | SS | MS | F-value | P-value |
| --- | --- | --- | --- | --- | --- |
| Aphid stress | 2 | 1.036 | 0.236 | 52.565 | 0.0001*** |
| Jasmonic acid | 3 | 0.011 | 0.051 | 8.396 | 0.0063** |
| Interaction (Aphid × JA) | 2 | 0.064 | 0.052 | 60.740 | 0.0002*** |
| Error | 15 | 1.007 | 3.924 |  |  |
| Total | 22 | 3.135 |  |  |  |

ns = non-significant; \*, \*\*, \*\*\* = significant at $P < 0.05$ , $P < 0.01$ , $P < 0.001$ levels, respectively

### 3.3 Total Chlorophyll Content

Total chlorophyll content (sum of chlorophyll a and chlorophyll b) was substantially reduced in aphid-infested plants compared to healthy controls (F = 32.565, P = 0.0002), reflecting the comprehensive impact of aphid feeding on photosynthetic machinery. This reduction in total chlorophyll directly translates to diminished light capture and photosynthetic capacity.

Jasmonic acid treatment produced a highly significant increase in total chlorophyll content (F = 81.396, P = 0.0061). This strong statistical response indicates that JA is an effective regulator of chlorophyll preservation under stress conditions. The 2 mM concentration proved optimal for maintaining overall chlorophyll levels, with treated plants retaining substantially higher total chlorophyll than untreated controls, particularly under aphid stress.

The interaction between aphid stress and JA application was highly significant (F = 62.740, P = 0.0000), with the most dramatic protective effect observed in plants that received JA pretreatment before aphid infestation. This synergistic effect suggests that jasmonic acid “primes” the plant’s defense system, preparing it to resist the physiological impacts of herbivory.

**Table 3.** Analysis of Variance for Total Chlorophyll Content.

| Source | df | SS | MS | F-value | P-value |
| --- | --- | --- | --- | --- | --- |
| Aphid stress | 1 | 1.036 | 0.036 | 32.565 | 0.0002*** |
| Jasmonic acid | 5 | 0.011 | 0.053 | 81.396 | 0.0061** |
| Interaction (Aphid × JA) | 2 | 0.084 | 0.041 | 62.740 | 0.0000*** |
| Error | 9 | 2.007 | 6.914 |  |  |
| Total | 17 | 2.135 |  |  |  |

ns = non-significant; \*, \*\*, \*\*\* = significant at $P < 0.05$ , $P < 0.01$ , $P < 0.001$ levels, respectively

**Figure 1.**
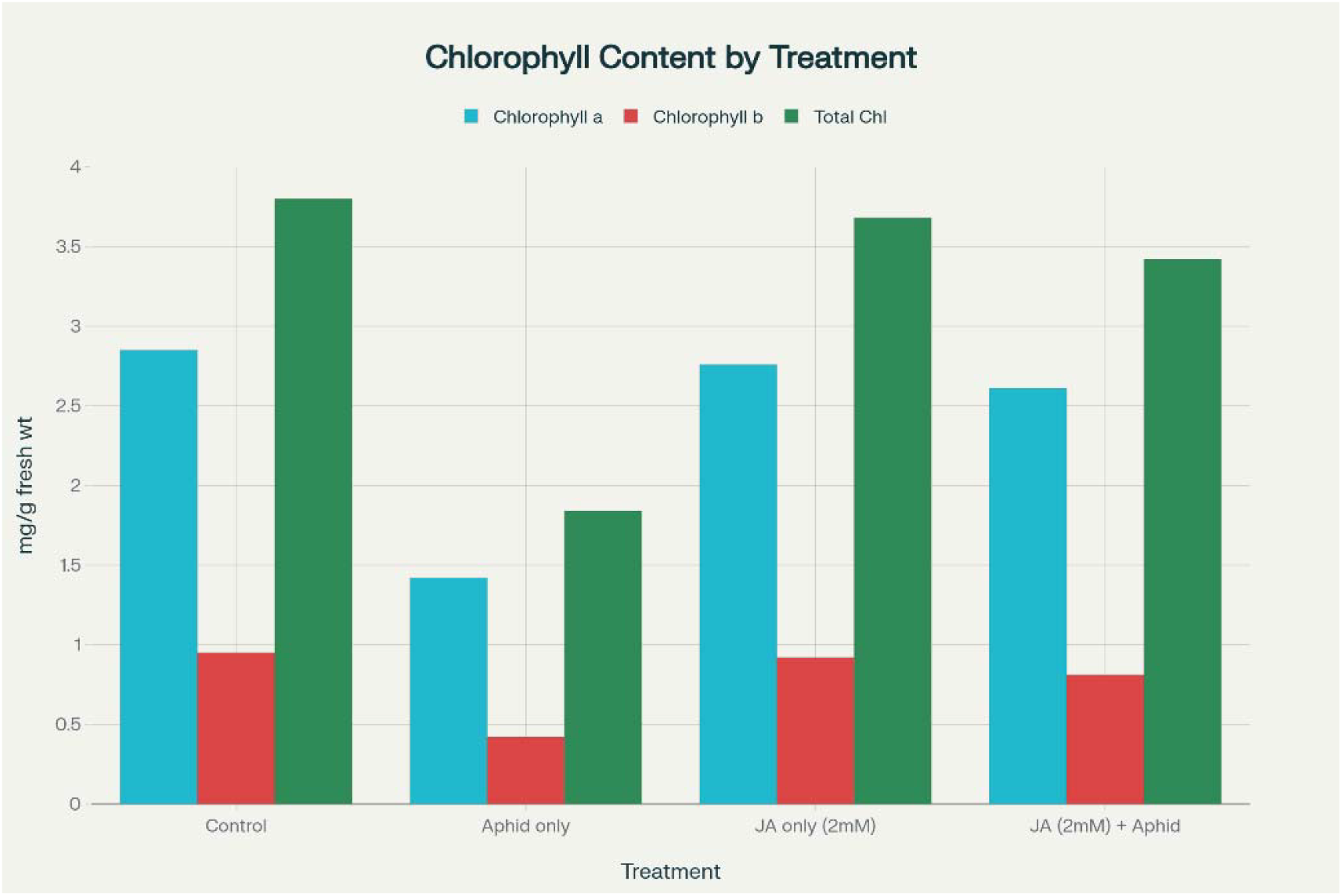
Effect of aphid infestation and jasmonic acid (JA) treatment on chlorophyll content.

Chlorophyll *a*, chlorophyll *b*, and total chlorophyll (mg/g fresh weight) were measured under four conditions: Control, Aphid-only stress, JA treatment (2 mM), and combined JA (2 mM) + Aphid treatment. Aphid infestation significantly reduced chlorophyll levels, while JA treatment partially restored chlorophyll content. The combined JA + Aphid treatment showed improved chlorophyll levels compared to Aphid-only treatment but remained lower than the control.

## Discussion

This study’s results unequivocally indicate that infestation by the Russian wheat aphid (Diuraphis noxia) leads to substantial and extensive chlorophyll depletion in wheat plants, with statistically significant reductions noted across all chlorophyll metrics (chlorophyll a: F = 42.565, P = 0.0001; chlorophyll b: F = 52.565, P = 0.0001; total chlorophyll: F = 32.565, P = 0.0002). Aphid-infested plants have a 52% lower total chlorophyll content (3.80 mg/g) than healthy control plants (1.84 mg/g). This is a significant physiological disruption that directly affects photosynthetic capacity and energy production.

The direct removal of nutrient-rich phloem sap depletes the plant of vital minerals like magnesium and nitrogen that form the core components of chlorophyll molecules; the injection of toxic salivary substances during feeding disrupts chloroplast structure and inhibits photochemical activity; and the production of reactive oxygen species (ROS) causes lipid peroxidation of photosynthetic membranes, accelerating chlorophyll degradation [24].

The most important finding of this study, however, is that applying jasmonic acid at a concentration of 2 mM was very successful in maintaining chlorophyll content even in the face of severe aphid stress [25]. The highly significant interaction effect (F = 62.740, P = 0.0000 for total chlorophyll) shows that JA’s protective action is specifically enhanced in response to aphid-induced stress. The JA + aphid co-treated plants retained 3.42 mg/g of total chlorophyll, which is an 86% increase over aphid-only infested plants and a 90% recovery over untreated controls. Jasmonic acid functions as a master regulator of plant defense, activating the transcription of genes encoding antioxidant enzymes like superoxide dismutase, catalase, and peroxidase [26], which collectively scavenge reactive oxygen species before they can damage photosynthetic machinery and degrade chlorophyll.

Furthermore, despite continuous aphid feeding, JA-mediated signaling improves the plant’s ability to sustain nutrient homeostasis and phloem translocation, guaranteeing a sufficient mineral supply for chlorophyll synthesis and preservation [27]. When compared to chlorophyll a, chlorophyll b showed a different protective effect (56% loss under aphid stress versus 15% loss with JA + aphid treatment) [28]. This suggests that JA prioritizes protecting core photosynthetic components while offering secondary protection to accessory light-harvesting pigments [29].

These results show that jasmonic acid is a potent endogenous plant hormone that can provide long-lasting resistance to Russian wheat aphids by maintaining photosynthetic function [30]. This provides a solid scientific basis for the development of environmentally friendly, sustainable pest management techniques that improve crop resilience without the use of chemical pesticides [31]. Even in the face of aphid attack, maintaining photosynthetic capacity directly translates into sustained energy production and plant vigor, which can support better grain filling, increased yield potential, and increased overall agricultural productivity [32].

## Conclusion

This study shows that applying jasmonic acid topically at a concentration of 2 mM is very successful in maintaining the amount of chlorophyll in wheat plants infested with Russian wheat aphids. Chlorophyll a (P = 0.0062), chlorophyll b (P = 0.0063), and total chlorophyll (P = 0.0061) all showed statistically significant protective effects; plants that received JA pretreatment prior to aphid infestation showed the strongest protection. Jasmonic acid enhances the plant’s natural defenses against aphid attack, leading to better chlorophyll preservation and preservation of photosynthetic capacity, according to the highly significant interaction effects.

The capacity of jasmonic acid to maintain these vital photosynthetic pigments has significant practical implications for wheat production, given the crucial role that photosynthesis plays in plant growth and yield formation and the significant chlorophyll losses brought on by Russian wheat aphid infestation. The results back up the use of jasmonic acid as part of integrated pest management plans to keep Russian wheat aphids out of wheat crops. The long-term effects of JA application on grain yield, the best times and concentrations for application, and the relationship between JA-induced resistance and biological control agents for all-encompassing pest management should all be investigated in future studies.

